# The pangenome of *Aspergillus fumigatus* highlights the dynamics of gene gain-loss over evolutionary timescales in a human fungal pathogen

**DOI:** 10.64898/2026.04.14.718449

**Authors:** Harry Chown, Johanna Rhodes, Matthew C. Fisher, Michael J. Bromley

## Abstract

How fungal pathogens generate and maintain genetic diversity under sustained environmental drug pressure is central to understanding the emergence of antifungal resistance. In environmental moulds such as *Aspergillus fumigatus*, escalating exposure to agricultural fungicides that share targets with clinical azoles imposes chronic selection outside the host, yet the genomic mechanisms enabling long-term adaptation remains unclear. Here, we reconstruct the largest eukaryotic pangenome assembled to date, comprising over 1,000 *A. fumigatus* isolates collected across 34 countries spanning a century. Using network-based orthogroup clustering combined with ancestral state reconstruction, we show that the *A. fumigatus* pangenome is open and shaped by continual gene gain and loss. Pangenome-wide association analyses identify accessory genes associated with itraconazole resistance, indicating that resistance evolution occurs within broader genomic backgrounds and beyond canonical target-site mutations. We further reveal that the accessory genome is structured into distinct evolutionary cohorts, including lineage-restricted gene sets enriched for mobile genetic element– associated domains, notably *Starship*-linked genes. These patterns suggest that *Starships* contribute to clade-specific genome architecture while remaining largely constrained by phylogenetic boundaries. Time-calibrated phylogenetic modelling reveals a relatively slow rate of gene turnover—approximately two orthogroup events per century—demonstrating that large-scale genome evolution in *A. fumigatus* is decoupled from elevated point mutation rates and contrasts sharply with bacterial systems. Together, these findings establish a quantitative framework for fungal pangenome evolution and reveal how structured accessory genome dynamics underpin antifungal resistance and long-term adaptation in this major human pathogen.

**Significance Statement:** Antifungal resistance in *Aspergillus fumigatus* threatens global health, yet the genomic processes enabling long-term adaptation under environmental drug pressure remain poorly understood. Using a global, century-spanning pangenome, incorporating over 1,000 isolates, we quantify gene gain and loss dynamics in this major human pathogen. We show that the pangenome is open but evolves slowly, with only ∼2 gene turnover events per century. Accessory genes follow two distinct evolutionary modes: some are lineage-restricted, while others are broadly distributed and dynamically exchanged. Resistance is associated with lineage-specific accessory genes, highlighting a broader role of resistance formation than target-site mutations alone. These findings reveal how heterogeneous turnover dynamics shape adaptation to widespread antifungal exposure.

## Introduction

*Aspergillus fumigatus* is an opportunistic human fungal pathogen and is the main aetiological agent of aspergillosis, a disease attributable to the mortality of an estimated 1.5 million people per year (1). The population is highly heterogeneous at the genomic and phenotypic level giving rise to variation in virulence and the development of fungal antimicrobial resistance (fAMR) (2–5). The frontline antifungal drug class are the azoles, which inhibits ergosterol biosynthesis, an essential cell membrane component (6). Resistance towards the azoles in *A. fumigatus* has been observed worldwide, with a background prevalence of ∼5% being observed in airborne spores (7, 8). The main resistance mechanism is a tandem repeat of the promoter region and non-synonymous nucleotide polymorphism in the gene encoding *cyp51A*, the azole target (TR_34_/L98H) (9).

Population genomics has revealed that the TR_34_/L98H genotype is strongly associated with population structure, with Cluster 3 (formerly known as Clade A) being the predominantly resistant lineage (10, 11). Additionally, isolates harbouring TR_34_/L98H genotypes have increased mutation rates, due to variants in the DNA mismatch repair system (MMR), enabling an enhanced propensity to develop fAMR to next-generation antifungals (12). Although the TR_34_/L98H genotype is well-documented, approximately 50% of clinical azole resistance cannot be accounted for by the presence of the TR_34_/L98H genotype alone (13). Previously, we have also identified gene presence-absence variants associated with itraconazole resistance (10, 11). Presence-absence variation has been identified through the generation of a pangenome – the total gene pool of a population - for *A. fumigatus*. Several pangenomes for *A. fumigatus* have been previously constructed, utilising a range of bioinformatic tools and collections of isolates (10, 11, 14–17). Genes within the pangenome can be characterised by their frequency within the population, with core genes found in the majority of the population, accessory genes conferring presence-absence variation, and singletons found only in a single isolate. Across studies on *A. fumigatus* there is a lack of concordance in pangenome statistics.

Pangenome size has been found to range from 10,005-15,309 orthogroups. The ratio of core, accessory and singleton orthogroups has been shown to vary with 41-78% of the gene pool comprising core genes. Additionally, it is common to describe a pangenome based on the concept of “openness” (18, 19). In an “open” pangenome the species gene pool is continually expanding, and the accessory genome is diverse. In contrast, a “closed” pangenome is characterised by the accessory genome reaching saturation – such that few or no novels genes are expected to be discovered. Openness can be calculated using gene accumulation curves fitted with Heaps’ power laws to measure the diminishing number of new genes with each additional genome (19). Currently, there is a lack of agreement across studies as to whether the pangenome is open in *A. fumigatus*, using the accumulation curve method (10, 14–16).

The mechanisms of gene gain and loss can vary across the Kingdom Fungi. Genomic regions can be gained through processes of duplication, sexual recombination and activity of mobile genetic elements (MGEs). In *A. fumigatus*, bioinformatic analyses have provided evidence for horizontal gene transfer (HGT) using both composition-based methods, which detect atypical base usage, and phyletic distribution approaches (20–22). Recently, a novel class of MGEs, called “*Starships*”, have been shown to actively transpose from *Paecilomyces variotii* into *A. fumigatus* (23). *Starship* elements are large MGEs, typically spanning ∼20-700 kilobases, that encode a “captain” tyrosine recombinase containing a DUF3435 domain thought to mediate their mobilization. Structurally, *Starships* are flanked by target-site duplications, with the captain gene positioned at the element’s 5′ end, followed by a suite of cargo genes that are co-transposed during mobilization (24). In *A. fumigatus, Starships* carry genes that enhance spore survival under heat and UV stress, modulate virulence, support low oxygen growth and are associated with biofilm formation (25). Although there is growing evidence that *Starships* influence clinically relevant phenotypes, their transposition rate and population-level dynamics remain unknown in *A. fumigatus* (25). Understanding the activity of such mobile elements is particularly important given that other sources of gene flow, such as sexual recombination, appear to be rare in this species (26). This suggests that transposition and related mobile genetic processes may represent alternative mechanisms shaping genomic diversity and adaptation. Quantifying gene gain-loss is therefore pivotal to understanding how *A. fumigatus* adapts to selective pressures imposed by antifungal drugs and agricultural fungicides.

We sought to quantify gene gain-loss across both temporal and evolutionary timescales using the pangenome of *A. fumigatus*, enabling assessment of pangenome openness and the rate at which resistance-associated genes are acquired. We show that the overall rate of pangenome evolution is slow and largely consistent across phylogenetic backgrounds, although distinct cohorts of accessory genes undergo accelerated gain-loss dynamics within the population. *Starship* captains display strong lineage dependence indicating constrained mobility across clusters. Together, these results reveal fine-scale plasticity within an otherwise stable pangenome and demonstrate how structured accessory genome dynamics shape adaptation in this mould.

## Results

### The pangenome of *Aspergillus fumigatus* is continuously expanding

Previous pangenomic studies in *A. fumigatus* have utilised different gene clustering techniques to construct a pangenome for the species (10, 14–17). In our initial study on 1,098 globally-acquired *A. fumigatus* isolates, we noticed that paralogous genes, such as *cyp51A* and *cyp51B* were incorrectly clustered together, as observed in other studies (16). To improve the resolution and investigate the effect of gene clustering on the pangenome, we first re-clustered the pangenome of *A. fumigatus* using a higher stringency network-based approach and compared this to the initial dataset (Figure 1) (Dataset S1). Utilisation of the network-based method resulted in a doubling of pangenome size from the initial 10,005 orthogroups to 22,198. The relative distribution of Core:Accessory:Singleton orthogroups remained consistent between the employed techniques (Network: 42:27:31; OrthoFinder: 41:29:30; Figure 1. B,C). However, comparisons of orthogroup accumulation curves revealed a change in the degree of pangenome “openness” – the potential for new genes to be identified with increased sampling of the population (Figure 1. D). The network-approach identified higher degrees of pangenome openness, as calculated by α-statistic (Network = 0.60, OrthoFinder = 0.83). This difference was also observed after the removal of singletons from the calculation (Network = 0.71 and OrthoFinder = 1.00), with the OrthoFinder method indicating a closed pangenome when there is no consideration of singletons.

**Figure 1.**
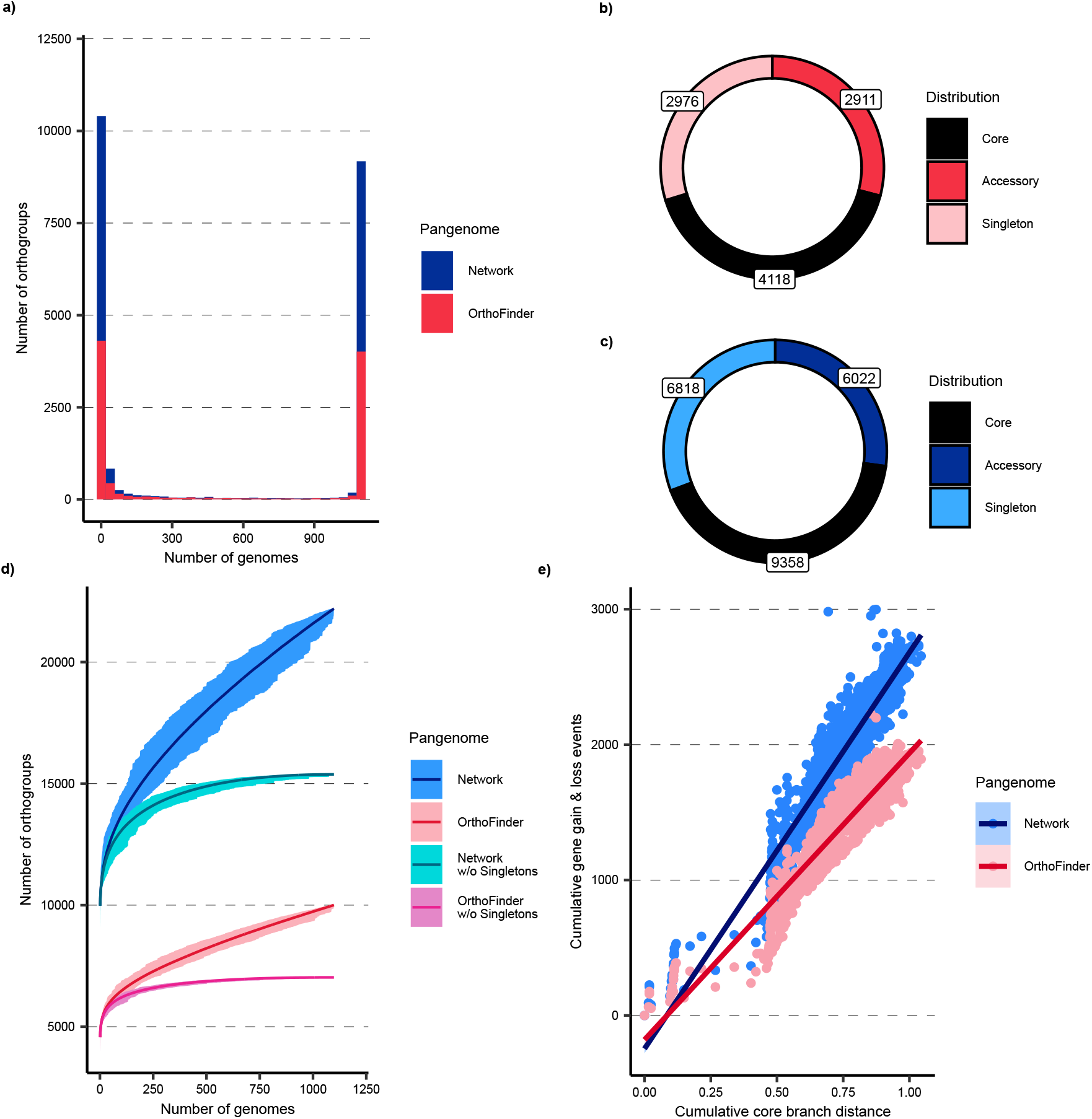
The “open” pangenome of *A. fumigatus* can fluctuate in pangenome size and openness estimation when using different orthogroup clustering techniques. **a**, Histogram of orthogroup distribution, using two pangenome clustering techniques: the protein-identity network method, using BLASTP and NetworkX (blue), and OrthoFinder (red). **b**, Donut plot summarising the total number of core (black), accessory (dark blue) and singleton (light blue) orthogroups calculated using the network method. Actual counts for each group are displayed in text boxes. **c**, Donut plot summarising the total number of core (black), accessory (dark red) and singleton (light red) orthogroups calculated using OrthoFinder. Actual counts for each group are displayed in text boxes. **d**, Orthogroup accumulation curves for network (dark blue) and OrthoFinder (dark red) methodologies and with singletons removed for each method (w/o) (light blue and light red). Rarefaction was carried out using 1,000 permutations. The mean across permutations is represented as a line and the lighter ribbon indicates the maxima and minima for each permutation. Heap’s Law was applied to each accumulation curve and the summary statistic depicting openness, α, was calculated (Network = 0.60, OrthoFinder = 0.83, Network w/o Singletons = 0.71, OrthoFinder w/o Singletons = 1.00). **e**, Cumulative orthogroup gain and loss events for each branch in the phylogeny, calculated through ASR for network and OrthoFinder methodologies, versus the cumulative branch lengths starting from the root in the species phylogeny. Each point represents a node in the phylogeny. Lines represents a linear model fit (Network: R^2^ = 0.888, p-value < 2.2 x 10^-16^; OrthoFinder: R^2^ = 0.854, p-value < 2.2 x 10^-16^).

Due to the variation in these results, we used Panstripe (27) to calculate pangenome openness through ancestral state reconstruction (ASR) of the gene pool at each node within the phylogeny. Openness is then assessed based on the cumulative number of orthogroup gain and loss events as the phylogenetic tree is traversed (cumulative branch length) (Figure 1. E). Similar to the gene accumulation curves, we found that both models were considered open and that the network methodology had higher inferred openness compared to OrthoFinder. This was shown through higher positive correlation between branch lengths and cumulative gain-loss (R^2^ values: Network=0.888; OrthoFinder=0.854) and observed through higher core statistics from Panstripe (Fig. S2).

These findings support an open model for the pangenome of *A. fumigatus*; thus the accessory gene pool is undergoing continuous expansion. Although gene clustering techniques can affect pangenome estimates, openness can be inferred through novel methods that relate pangenome size to the phylogeny.

### Accessory orthogroups are associated with azole resistance and phylogenetic cluster

Previously we have shown that accessory genes are associated with drug resistance and phylogenetic structure using pangenome-wide association studies (panGWAS) (10, 11). Therefore, with the higher resolution network-based pangenome, we expected to uncover additional resistance-associated orthogroups. Using a subset of isolates with known itraconazole susceptibility-profiles, identified in the previous study (*n*=319) (11), we found 85 accessory orthogroups that are associated with itraconazole resistance (Dataset S2). Again, the majority (*n*=84) were overrepresented in phylogenetic Cluster 3, the lineage containing the highest proportion of azole resistant genotypes (Fig. 2A). These findings suggest that resistance-linked accessory genes are largely confined to specific evolutionary lineages.

**Figure 2.**
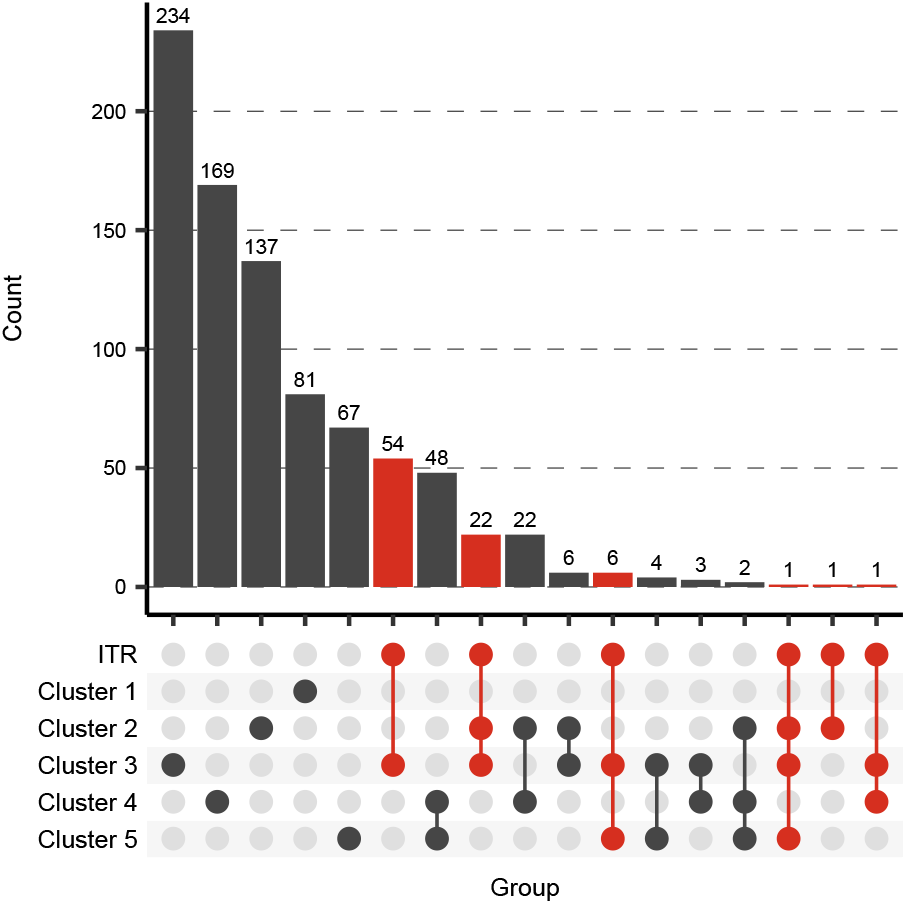
Pangenome-wide association studies reveals accessory orthogroups that are associated with itraconazole resistance and phylogenetic clusters. Upset plot showing the intersection of statistically associated accessory orthogroups across population clusters (Clusters 1-5) and itraconazole resistance (ITR). The bar chart at the top represents the number of shared accessory genes for each combination of cluster and ITR associations. Dots and connecting lines underneath illustrate the specific intersections between clusters and resistance phenotypes, highlighting commonly associated orthogroups. ITR associations are coloured in red.

Annotation analysis revealed that approximately half of the resistance-associated orthogroups had identifiable protein domains (*n*=38). Two orthogroups likely arose from bacteriophage contamination by a PhiX sequencing control. Nine were present in the *A. fumigatus* Af293 reference genome. The predicted functions of genes not found in the reference genome spanned enzymatic activity, transcriptional regulation, signal transduction and secondary metabolite biosynthesis. Several orthogroups encoded domains implicated in protein-protein interactions, including NB-ARC, NACHT, ankyrin repeats, and F-box-like motifs. One orthogroup encoded ABC transporter domains and several others contained domains of unknown function (e.g., DUF3435) (Dataset S2).

Beyond resistance, panGWAS identified many orthogroups associated with population structure (Fig. 2A). The average number of cluster-associated orthogroups was 137 per cluster and Cluster 3 contained the highest number of overrepresented orthogroups (*n*=234). These findings reinforce the strong relationship between accessory genome content and phylogenetic lineage in *A. fumigatus* and show that many azole-resistance associated genes are phylogenetically stratified.

### Temporal analysis of the pangenome indicates a slow rate of orthogroup acquisition and loss

Recent findings of elevated mutation rates in certain *A. fumigatus* sub-populations have raised the hypothesis that specific lineages may be evolving more rapidly than others (12). However, the tempo of gene gain and loss across natural populations – especially in real time – remains poorly characterised. To address this, we estimated orthogroup turnover rates in real-time across the *A. fumigatus* population and tested whether these rates vary between phylogenetic clusters.

Using a time-calibrated phylogeny from our previous study comprising of 512 isolates (11), alongside the corresponding binary network-based pangenome matrix, generated in this study, we were able to model orthogroup gain and loss across the dated tree. Similar to the comparison against the SNP-based phylogeny, ASR was performed to infer orthogroup presence or absence along branches, allowing us to estimate orthogroup gain and loss events both at the whole-population level and within individual phylogenetic clusters.

We observed a consistent accumulation of orthogroup gain and loss events over time (Fig. 3A), with an estimated rate of approximately two orthogroup turnover events (gain or loss) per 100 years (Fig. 3B). Notably, the rate of orthogroup exchange appeared relatively uniform across most clusters, although Cluster 1 exhibited the highest variability in turnover rate (Fig. 3B). These results suggest that while some variability exists, the average rate of orthogroup birth-death processes in *A. fumigatus* across all orthogroups is low and stable over time.

**Figure 3.**
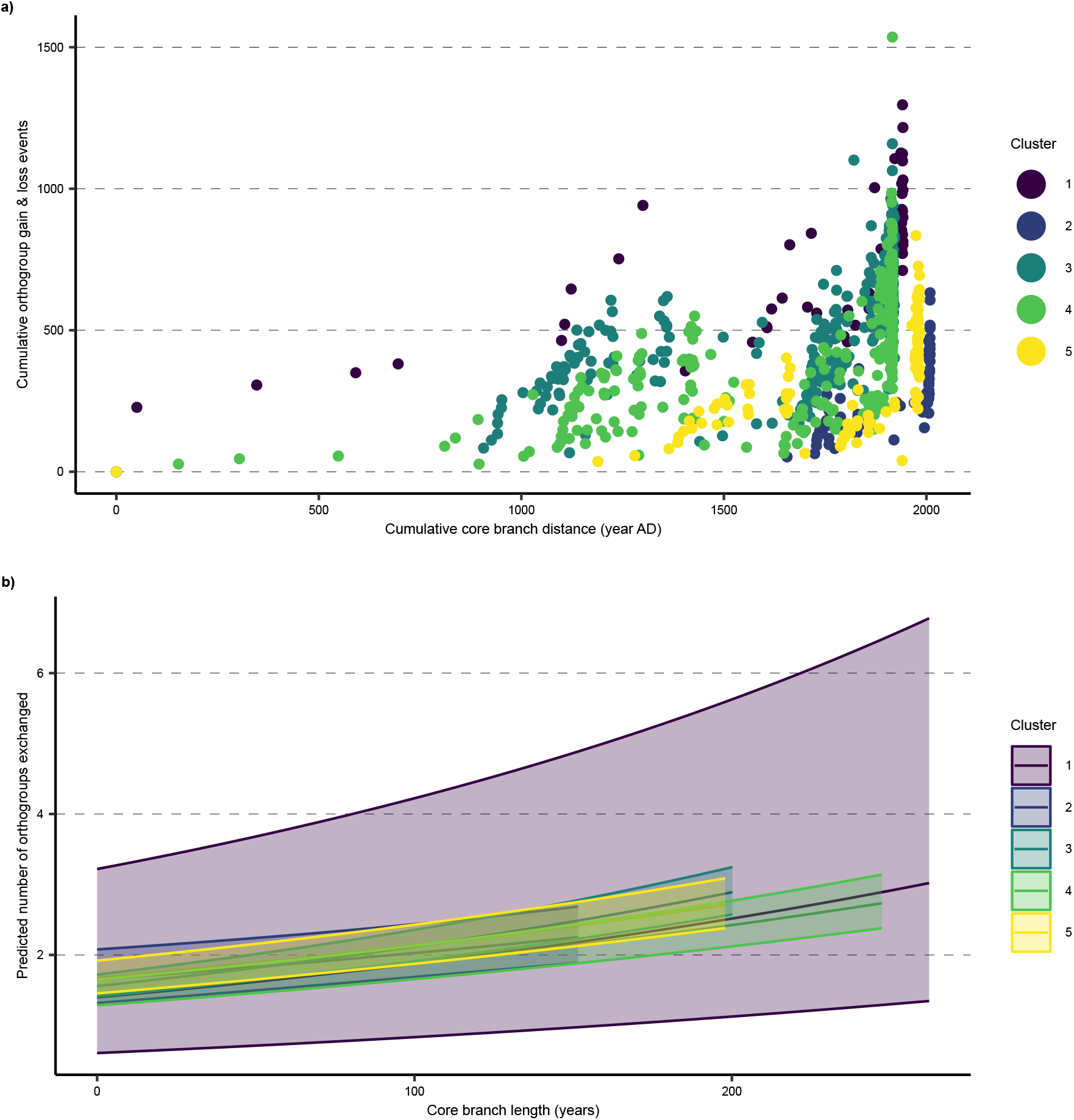
Slow rate of total orthogroup birth-death across lineages identified through temporal analysis. **a**, Cumulative orthogroup gain and loss events for each branch in the time-calibrated phylogeny, calculated through ancestral state reconstruction, versus the cumulative branch length, in years, starting from the root in the species maximum clade credibility tree. Each point represents a node in the phylogeny. Colours represent phylogenetic clusters (purple=1, dark blue=2, dark green=3, green=4, yellow=5). **b**, Relationship between core branch length, in years, and the predicted number of gene birth-death events per cluster and in total (red). Accompanying ribbon illustrates the 95% confidence interval of each model’s fit per lineage.

### Accessory orthogroups undergo differential rates of gain and loss over evolutionary timescales

Gene presence-absence variation in *A. fumigatus* can arise through multiple mechanisms, including transposition, HGT, gene duplication and mutagenesis. MGEs - such as transposons and *Starships* - are particularly dependent on transposition and HGT for their mobility within populations (23, 28, 29). As observed in bacterial systems, distinct MGE classes exhibit unique gain and loss profiles within pangenomes (30). However, this phenomenon will be masked when analysing total gene turnover as over 40% of the pangenome is core. Therefore, we characterised gain and loss patterns across individual orthogroups to uncover the heterogenous dynamics of accessory orthogroups. We first assessed the phylogenetic distribution of accessory orthogroups, present in 1-99% of the population, using the Fritz and Purvis’ D statistic (31) (Dataset S4). Of the 3,304 accessory orthogroups, 1,485 exhibited lineage-dependent distributions (D=0) (Fig. 4A), whereas the remaining 1,819 varied independently of the phylogeny, with a subset of these (*n*=589) indistinguishable from random expectations (D=1) (Fig. 4A). GAM analyses confirmed that orthogroups with D=0 displayed decreasing similarity in presence-absence profiles with increasing phylogenetic distance, consistent with evolutionary inheritance, whereas orthogroups with D=0 showed attenuated or absent phylogenetic structure (Fig. 4B). These patterns indicate that accessory orthogroups undergo gain and loss at heterogeneous evolutionary rates relative to the phylogeny and that closely related isolates may differ substantially in their accessory gene repertoires.

**Figure 4.**
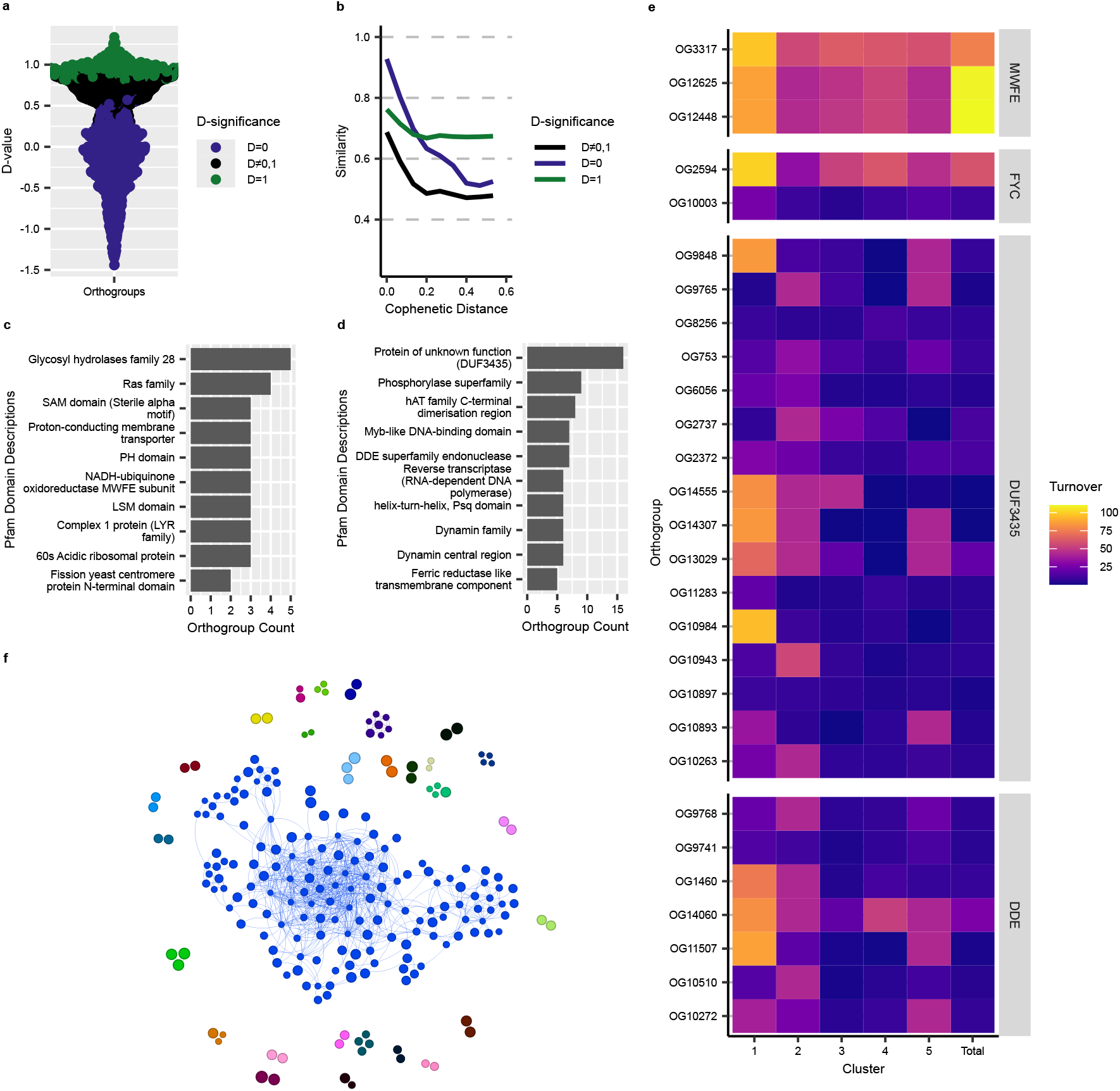
Heterogeneous evolutionary behaviour of accessory orthogroups. **a**, Distribution of Fritz and Purvis’ D statistic for all accessory orthogroups. Colours represent values significantly equal to 0 (blue), significantly equal to 1 (green) or neither (black). **b**, Generalised additive model (GAM) fits describing the relationship between cophenetic (pairwise phylogenetic) distance and Jaccard similarity of presence-absence profiles for orthogroups assigned to the three D-significance categories. **c**, Pfam domain descriptions occurring exclusively in orthogroups with D=0, showing the ten domains represented by the largest number of orthogroups. **d**, Pfam domain descriptions occurring exclusively in orthogroups with D=0, showing the ten domains represented by the largest number of orthogroups. **e**, Orthogroup turnover rates for selected Pfam domains: NADH-oxidoreductase MWFE subunit (Pfam: PF15879; MWFE), Fission yeast centromere protein N-terminal domain (Pfam: PF18107; FYC), DUF3435 domains (Pfam: PF11917) and DDE superfamily endonucleases (Pfam: PF13358; DDE). Turnover is shown per cluster and across the full population. **f**, Co-occurrence network of orthogroups with D=0. Nodes represent orthogroups; edges indicate co-occurrence; node size scales with D statistic; colours denote network components.

To explore possible functional biases, we compared Pfam domain content between lineage-dependent and lineage-independent groups. Although no domain class was significantly enriched, several domains were found exclusively within cohorts. The ten most represented lineage-dependent domains were mostly associated with MGEs including *Starship* captains (DUF3435), hAT-family transposases (hAT dimerization, DDE endonuclease, Psq/HTH, Myb-like DNA-binding) and retrotransposon-derived proteins (reverse transcriptase). A smaller number encoded dynamin-family GTPases involved in membrane remodelling and one domain was a ferric reductase-like transmembrane domain associated with iron acquisition (Fig. 4C). In the lineage-independent cohort, domain analysis revealed a more functionally diverse set, including components of the mitochondrial Complex I (LYR and MWFE), RNA processing (LSM domain and 60S acidic ribosomal protein) and cellular signalling (Ras GTPase, PH and SAM domains). Additional domains were linked to extracellular carbohydrate degradation (GH28) and chromosome segregation (centromere protein N-terminal domain) (Fig. 4D).

We sought to characterise the DUF3435 domains as *Starship* captains and investigated whether the phylogenetic signal is only observed in specific *Starship* families. We were able to characterise 15 out of 16 lineage-dependent orthogroups containing DUF3435 domains as *Starship* captains. The *Starship* captains were assigned to one of five families, including Enterprise (n=3), Galactica (n=2), Hephaestus (n=1), Prometheus (n=5) and Tardis (n=4) (Dataset S3). This highlights that a diverse range of *Starship* captains exhibit strong phylogenetic signal with limited shared presence between lineages.

As roughly half of the accessory orthogroups displayed clustering in a phylogenetic manner, we next examined whether their gain-loss dynamics varied across lineages. To provide an overview, we first characterised the rates of gain and loss relative to the phylogeny for orthogroups containing the most represented lineage-dependent and lineage-independent domains (Fig. S3). We found that orthogroups containing the DUF3435, DDE superfamily endonuclease, MWFE and fission yeast centromere protein domains observed the greatest diversity in inferred gain and loss rates (Fig. S3). Turnover rates (combined gain and loss) revealed that lineage-independent orthogroups exhibited higher turnover consistently across clusters, whereas lineage-dependent orthogroups varied markedly between lineages (Fig. 4E). Cluster 1 displayed the highest turnover rates for multiple domains, consistent with the elevated evolutionary flux as seen previously in this lineage (Fig. 3B).

Finally, to assess whether orthogroups influence each other’s distributions, we constructed a co-occurrence network for lineage-independent orthogroups. The network comprised of 29 connected components, including one large component of 151 orthogroups (Fig. 4F). Eleven hub orthogroups showed high connectivity (32-50 co-occurrences), although only three contained characterised Pfam domains (phosphotransferases, PalI-pH pathway, serine hydrolases). Additional highly connected orthogroups included those containing ankyrin repeats, WD domains and combined centromere-associated and transposase-associated domains (Dataset S4). These hubs may represent genetic contexts that promote the retention or acquisition of other accessory genes.

Together, these analyses reveal that the accessory gene pool of *A. fumigatus* undergoes heterogenous evolutionary processes. Orthogroups are either strongly structured by phylogeny or nearly randomly distributed across the population, with markedly different turnover behaviours. Moreover, co-occurrence patterns suggests that subsets of lineage-independent orthogroups form tightly connected modules that may facilitate additional gene acquisitions. These findings reveal the complex and dynamic architecture of the *A. fumigatus* pangenome.

## Discussion

In this study, we set out to quantify how gene gain and loss shape the long-term evolution of *Aspergillus fumigatus* across a period of increasing environmental drug pressure. By reconstructing a large, time-calibrated pangenome spanning more than a century of global sampling, we move beyond static descriptions of accessory gene content to directly measure the tempo and structure of pangenome evolution.

Our analyses reveal the dynamics of gene gain and loss in the pangenome of *A. fumigatus* and demonstrate that estimates of pangenome size and openness are strongly influenced by gene clustering methodology. By applying a stringent network-based clustering approach and integrating ASR, we confirm that the *A. fumigatus* pangenome is open, with continual acquisition of novel genes as additional isolates are incorporated. We further identify accessory orthogroups associated with itraconazole resistance and show that orthogroup turnover rates vary markedly across functional categories. Together these findings advance understanding of fungal genome plasticity and clarify how structured pangenome dynamics contribute to adaptive evolution under antifungal selection.

Pangenomic re-analysis revealed that the total number of orthogroups in *A. fumigatus* nearly doubled when network-based clustering was applied compared with OrthoFinder, while the ratio of core to accessory genes remained constant. This expansion largely reflects improved discrimination of paralogous genes, such as *cyp51A* and *cyp51B*, which were incorrectly merged in earlier analyses (11, 16). The stability of the core-to-accessory ratio despite increased orthogroup resolution indicates that most newly resolved gene families represent divergent homologues already present across the species, rather than genes uniquely present in subsets of isolates. These results suggests that gene divergence in *A. fumigatus* is dominated by vertical evolutionary processes rather than sporadic acquisition of highly divergent genes. This pattern contrasts sharply with bacterial pangenomes, where frequent HGT can introduce sequences with little or no homology to the existing gene pool; as a result, clustering thresholds strongly influence the apparent size of the accessory genome in Bacteria (32).

The openness of the *A. fumigatus* pangenome has been reported inconsistently across previous studies, largely reflecting differences in methodology, and sample diversity (10, 14–16). Despite methodological differences between the clustering approaches used here, both supported an open pangenome model. However, gene accumulation curves do not account for population structure and underlying errors in the presence and absence of genes which can inflate accessory gene values (33). ASR offers a robust approach, which investigates the birth-death rate of orthogroups whilst filtering out low abundance genes (33). Application of an ASR method to our dataset (27) further substantiated the conclusion that the *A. fumigatus* pangenome is open, revealing a strong linear relationship between cumulative orthogroup gain-loss and evolutionary distance. Pangenomic models used in this analysis represent binary representations of orthogroups showing that gene duplication alone is not a contributing factor of openness estimation, however gene duplication followed by diversification is the likely common mechanism of orthogroup birth. Importantly, pangenome openness of *A. fumigatus* likely reflects not only ongoing gene flux but also the large and heterogeneous genomic landscape of the species. As phylogenetically diverse isolates were incorporated, diversity increased across both accessory gene content and core genomic variation, indicating that continued sampling captures previously unsampled accessory regions. This pattern is consistent with a species characterised by a large effective population size and broad geographical and ecological distribution. These findings also highlight a fundamental challenge in fungal pangenomics: the absence of standardized criteria for orthogroup inference and analysis. Harmonizing clustering parameters and benchmarking against manually curated ortholog sets will be essential to ensure future reproducibility and cross-study comparability.

Our pangenome-wide association study (panGWAS) identified 85 accessory orthogroups significantly associated with itraconazole resistance, the majority of which were also overrepresented within phylogenetic Cluster 3, the lineage encompassing most azole-resistant isolates. The co-association between lineage-specific accessory gene content and itraconazole resistant associated gene content suggests that resistance evolution in *A. fumigatus* occurs within wider genomic backgrounds than mutations of target genes alone. This pattern is consistent with the co-selection of accessory loci that confer additional adaptive advantages under azole exposure or environmental stress. Such loci may functionally interact with resistance determinants, compensating for associated fitness costs, or enhancing survival in selective environments. The presence of ABC transporter domains within the itraconazole-associated set of accessory orthogroups represents a known mechanisms of azole efflux (34–37). In addition, the presence of orthogroups containing domains involved in signalling and protein–protein interaction (e.g., NACHT, NB-ARC, ankyrin repeats) hint at potential regulatory or stress response roles. However, functional interpretations remains challenging as nearly half of resistance associated orthogroups lack identifiable protein domains. Experimental validation will therefore be required to disentangle whether these accessory orthogroups directly contribute to antifungal resistance, buffer pleiotropic consequences of resistance mutations, or provide alternative adaptations in azole-containing environments.

Although the canonical resistance genotype (TR_34_/L98H) shows no detectable fitness defect in laboratory-constructed strains, naturally occurring azole-resistant isolates display variable fitness, reflecting broader genomic heterogeneity (38–40). Moreover, most prior studies have been conducted *in vitro* and fail to assess fitness within an ecological context in which accessory genes may confer additional advantages. The lineage specificity of resistance-associated orthogroups suggests maintenance of accessory genes by the same process which has been driving the emergence of the resistant subpopulation: local agricultural azole selection, clonal expansion via asexual recombination, and limited inter-lineage recombination (11).

In this study, we provide the first temporal estimate of gene gain-loss in *A. fumigatus*, revealing a relatively slow average turnover rate of approximately two orthogroup events per century. This modest rate contrasts sharply with bacterial systems, where pangenome turnover can occur orders of magnitude faster. The relative genomic stability of *A. fumigatus* likely reflects its predominantly clonal reproduction, limited HGT, and strong purifying selection maintaining core gene content. Notably, while the overall rate was low, cluster-specific variability was only observed within Cluster 1. In contrast, isolates from Cluster 3 —previously shown to exhibit elevated mutation rates due to defects in the DNA MMR pathway (12) — did not display higher gene turnover. This suggests that variation in point mutation rate does not directly translate into accelerated gene gain-loss, indicating that large-scale genomic divergence in *A. fumigatus* is decoupled from nucleotide-level mutational processes.

Comparative interpretation of turnover rates remains challenging due to the scarcity of equivalent temporal models in Kingdom Fungi. No previous studies have quantitatively assessed gene gain-loss across dated fungal phylogenies, making it difficult to determine whether the rate observed here is unusually low or broadly representative of filamentous fungi. For instance, the pangenome of *Aspergillus flavus* exhibits a similar number of core genes but has been described as closed, suggesting that gene gain-loss occur at lower rates relative to the more dynamic *A. fumigatus* population (41). Developing comparable pangenomic and phylogenomic frameworks across diverse fungal species are therefore essential in order to contextualize gene turnover dynamics within the broader landscape of fungal evolution.

Our evolutionary analyses revealed that the accessory genome is not homogenous in its behaviour but is instead partitioned into evolutionary distinct cohorts. Approximately half of all accessory orthogroups exhibit strong phylogenetic structure, implying lineage restricted inheritance, as previously identified in azole-resistance associated orthogroups. Phylogenetic structuring of accessory gene composition may facilitate niche adaptation in sub-populations, creating ecological divergence, and potentiate speciation events (42–44). This lineage dependent class is enriched for MGE domains, including *Starship* captains and putative transposons – suggesting that MGE distribution may be clade-specific. Although mobilisation of *Starships* in *A. fumigatus* has been observed (25), these results suggest that movement of *Starships* may be confined within phylogenetically conserved lineages and that the population has developed barriers towards HGT of *Starships* or the presence of strict selective constraints on *Starship* cargo genes. In contrast, the lineage-independent cohort comprises a diverse collection of co-occurring orthogroups. This implies that accessory genome dynamics involves the coordinated turnover of multi-gene modules. Although sexual reproduction is rare, recombination among diverse environmental isolates likely facilitates the redistribution and long-term maintenance of these accessory loci across lineages (26). Although co-occurrence can infer shared biological properties within cohorts of genes, the majority of co-occurring orthogroups do not have annotated protein domains (45). The mechanisms underlying these associations remain unresolved and synteny analyses using long-read-sequenced genomes, together with targeted functional investigations, will be essential to determine whether co-occurrence is driven by physical co-localisation, shared biological roles, or other evolutionary forces.

Together, our analyses reveal a mosaic accessory genome in *Aspergillus fumigatus* shaped by the interplay of vertical inheritance, gene duplication and diversification, structured gene flux, and lineage-specific mobile genetic elements. These population genomic processes underpin the long-term architecture of an expanding gene repertoire during a period of unprecedented escalation in agricultural fungicide use (46). By quantifying gene gain and loss across both contemporary and evolutionary timescales, this study provides the first dynamic, population-scale framework for understanding how the accessory genome of a major human fungal pathogen evolves in response to sustained environmental antifungal selection.

## Materials and Methods

### Initial pangenomic data

We analysed a previously generated pangenome comprising of 1,098 *A. fumigatus* genomes (11) described in detail in the original study. Briefly, whole-genome sequencing data from 1,102 isolates were collected across 34 countries and 102 years, spanning six continents, were quality-filtered, assembled *de novo* with SPAdes v3.15.4 (47), and scaffolded against the CEA10 reference genome (NC_017016.1) (48). Assemblies were retained based on a minimum 20x read coverage compared to the Af293 reference (GCF_000002655.1 ASM265v1) (49), species identity (Kraken2 v2.1.2 (50) read classification > 80%) and completeness (BUSCO v5.4.3 (51) >80%), yielding a final set of 1,098 genomes. Reference genes were detected with Exonerate v2.2.0 (52) and the Af293 reference proteome; novel genes were predicted, using BRAKER v3.0.2 (53) with the OrthoDB v11 dataset (54), in unannotated regions and putative contaminants were excluded based on low coverage and lack of *A. fumigatus* homology from DIAMOND BLASTP v2.1.8.162 (55) searches against a clustered NCBI protein database (56, 57). The pangenome was then constructed from all predicted protein sequences using OrthoFinder v2.5.5 (58), with matrices updated to include reference homologs.

An overview of the underlying dataset, including the reconstructed phylogeny and the distribution of isolates by geography and collected date is provided in Fig. S1. The phylogeny with corresponding lineage information was visualised in in R v4.2.0 (59) using the packages “ape” v5.7-1 (60), “ggtree” v3.16.3 (61) and “ggtreeExtra” v1.18.1 (62). Geographic distributions of samples and their collection date, if known, were visualised by obtaining a map using “giscoR” v1.0.0 (63), retrieving country coordinates using “sf” v1.0-23 (64, 65), adding pie-charts to the map with “scatterpie” v0.2.6 (66) and visualisation using “ggplot2” v4.0.1 (67) and “retroPal” v0.2.1 (68). Data was manipulated using the “dplyr” v1.1.4 (69) and “tidyr” v1.3.1 (70) packages.

### Pangenome reconstruction

Initial OrthoFinder analysis clustered true paralogs of *cyp51A* and *cyp51B* together indicating insufficient resolution (11, 16). To increase specificity, the pangenome was reconstructed using a network-based approach. Annotated protein sequences from all isolates were compared pairwise with BLASTP v2.15.0 (71), using a minimum percentage identity and query coverage of 90%. The resulting edge list was processed with a custom Python v.3.9.13 (72) script employing ‘networkx’ v2.2 (73) to generate orthogroups from unconnected network components. This procedure identified representative hubs and produced matrices of gene identities, copy numbers and binary presence-absence data across isolates. Storage and manipulation of data was carried out using the ‘pandas’ module.

### Domain annotation

A representative pangenome was generated for each orthogroup using the hub orthogroups from the network approach. The representative pangenome underwent protein domain classification using InterProScan v5.67-99.0 (74) with all databases.

### Pangenome association testing

Utilising the network-based pangenome, association tests were performed to identify associations between the presence of orthogroups and itraconazole resistance and cluster assignment. This was carried out using Scoary v1.6.16 (75), supplied with the corresponding SNP-based phylogeny. Analyses on resistance and clusters used a minimum *p*-value of 0.05. For resistance and cluster associations, *p*-values underwent Bonferroni correction.

### Accumulation curves

For each pangenome, the distribution of orthogroups amongst the population was then calculated and summarised, whereby a core orthogroup is present in >95% of isolates, a singleton is present in a single isolate, and the remainder are classed as accessory. To identify the “openness” of the pangenome and show the differences between pangenome construction methods, orthogroup accumulations curves were generated for both approaches. In both cases, singletons were also removed and visualised to show the effect of singletons on the study. Accumulation curves were ran with 1,000 permutations and the a decay value for pangenome openness was calculated, this was performed using “micropan” v2.1 (76) in R v4.2.0 (59).

### Pangenome evolution with Panstripe

Pangenome evolution was evaluated using Panstripe v0.2.0 (27). The network-based pangenome matrix was analysed in comparison to two previously generated (11): (i) reference-based tree including all isolates constructed from SNPs aligned to the Af293 genome (GCF_000002655.1 ASM265v1) (49), and (ii) a time-calibrated tree including isolates with known collection dates and itraconazole susceptibility profiles, derived from mitochondrial SNPs and maximum clade credibility tree estimation with BEAST2 v2.7.1 (77). All trees were re-rooted to the outgroup *Aspergillus neolipticus* isolate, C250, using “ape” v5.7-1 (60) in R.

Panstripe analyses were performed on the total population and individual phylogenetic clusters across both reference- and time-based trees, and for pangenomes constructed with OrthoFinder and the network approach. To account for skewed branch-length, a quasi-Poisson model was adopted. Panstripe was applied to calculate the relationship between branch length and cumulative orthogroup gain-loss events and between phylogenetic and pangenomic distance. Output data was visualised with “ggplot2” v3.4.4 (67).

### Quantifying pangenome divergence

To identify the impact of phylogeny on the distribution of orthogroups, phylogenetic signal of orthogroups was calculated using Fritz and Purvis’ D statistic within the “caper” v1.0.4 package (31, 78). This was performed only on accessory orthogroups that were present in more than 1% of the population and less than 99%. Two cohorts of orthogroups were identified based on the D statistic being either insignificantly (*p* > 0.05) or significantly different from zero. Domain enrichment, integrating the known Pfam domains, was performed using a Fisher’s exact test with Benjamin-Hochberg correction between groups.

We assessed how orthogroups vary in presence-absence over evolutionary timescales by comparing phylogenetic and pangenomic distances. Cophenetic distances were calculated from the reference-based phylogeny using the R package “ape”. Pangenomic distance was computed as pairwise Jaccard distance from the binary presence-absence matrix. Trends between pangenomic distance and phylogenetic distance were summarised by applying a generalised additive model (GAM) using “mgcv” v1.9-1 (79, 80) and “tidymv” v3.4.2 (81). This was carried out for cohorts of orthogroups whereby D=0, D=1 and D≠0,1 and for orthogroups containing Pfam domains only found in D=0 and D≠0 groups.

Orthogroups that contained DUF3435 domains (Pfam: PF11917), DDE superfamily endonucleases (Pfam: PF13358), Fission yeast centromere protein N-terminal domain (Pfam: PF18107) and NADH-ubiquinone oxidoreductase MWFE subunit (Pfam: PF15879) were selected for further analysis based on distinct GAM profiles. Turnover rates were calculated for these orthogroups using the M*k* model for discrete character evolution (82) with the All-Rates-Different (ARD) model from ‘phytools’ v2.5-2 (83).

To assess whether the presence of a lineage-independent orthogroup affected the presence of another, co-occurrence analysis was performed using Coinfinder v1.2.1 (84). Output from Coinfinder was then analysed in R, using ‘igraph’ v2.2.1 (85), and visualised in Gephi v0.10.1 (86).

### *Starship* captain family assignment

Representative protein sequences of phylogenetically dependent orthogroups containing DUF3435 domains underwent *Starship* captain classification based on similarity to HMM profiles of tyrosine recombinase (YR) reference sequences. YR reference sequences were obtained from Starfish v1.1.0 (87) and were aligned to DUF3435-domain containing proteins using hmmsearch v3.3.2 (88). A single family was assigned to each protein based on the lowest E-value.

## Acknowledgments

We would like to thank Dr. Nicholas Croucher and Dr. Maria Domingo-Sananes for their helpful discussions and suggestions for improving this study.

## References

1. D. W. Denning, Global incidence and mortality of severe fungal disease. Lancet Infect. Dis. 24, e428–e438 (2024).

2. F. Alshareef, G. D. Robson, Genetic and virulence variation in an environmental population of the opportunistic pathogen Aspergillus fumigatus. Microbiology 160, 742–751 (2014).

3. M. C. Fisher, et al., A one health roadmap towards understanding and mitigating emerging Fungal Antimicrobial Resistance: fAMR. Npj Antimicrob. Resist. 2, 36 (2024).

4. F. Puértolas-Balint, et al., Revealing the Virulence Potential of Clinical and Environmental Aspergillus fumigatus Isolates Using Whole-Genome Sequencing. Front. Microbiol. 10 (2019).

5. S. Zhao, W. Ge, A. Watanabe, J. R. Fortwendel, J. G. Gibbons, Genome-Wide Association for Itraconazole Sensitivity in Non-resistant Clinical Isolates of Aspergillus fumigatus. Front. Fungal Biol. 1, 617338 (2021).

6. H. Elsaman, et al., Toxic eburicol accumulation drives the antifungal activity of azoles against Aspergillus fumigatus. Nat. Commun. 15, 6312 (2024).

7. H. H. Kortenbosch, et al., Land use drives drug resistance in an airborne human fungal pathogen. ISME J. 19, wraf246 (2025).

8. J. M. G. Shelton, et al., Citizen science reveals landscape-scale exposures to multiazole-resistant Aspergillus fumigatus bioaerosols. Sci. Adv. 9, eadh8839 (2023).

9. F. Gsaller, et al., Sterol Biosynthesis and Azole Tolerance Is Governed by the Opposing Actions of SrbA and the CCAAT Binding Complex. PLOS Pathog. 12, e1005775 (2016).

10. J. Rhodes, et al., Population genomics confirms acquisition of drug-resistant Aspergillus fumigatus infection by humans from the environment. Nat. Microbiol. 7, 663–674 (2022).

11. M. Fisher, et al., Recent European origin of azole resistance in the critical priority fungal pathogen Aspergillus fumigatus. [Preprint] (2025). Available at: https://www.researchsquare.com/article/rs-7905776/v1 [Accessed 9 March 2026].

12. M. J. Bottery, et al., Elevated mutation rates in the multi-azole resistant Aspergillus fumigatus clade drives rapid evolution of antifungal resistance. [Preprint] (2023). Available at: https://www.biorxiv.org/content/10.1101/2023.12.05.570068v1 [Accessed 8 May 2024].

13. E. Snelders, et al., Emergence of azole resistance in Aspergillus fumigatus and spread of a single resistance mechanism. PLoS Med. 5, e219 (2008).

14. A. E. Barber, et al., Aspergillus fumigatus pan-genome analysis identifies genetic variants associated with human infection. Nat. Microbiol. 6, 1526–1536 (2021).

15. M. A. C. Horta, et al., Examination of Genome-Wide Ortholog Variation in Clinical and Environmental Isolates of the Fungal Pathogen Aspergillus fumigatus. mBio 13, e0151922 (2022).

16. L. A. Lofgren, B. S. Ross, R. A. Cramer, J. E. Stajich, The pan-genome of Aspergillus fumigatus provides a high-resolution view of its population structure revealing high levels of lineage-specific diversity driven by recombination. PLOS Biol. 20, e3001890 (2022).

17. C. G. P. McCarthy, D. A. Fitzpatrick, Pan-genome analyses of model fungal species. Microb. Genomics 5, e000243 (2019).

18. H. Tettelin, et al., Genome analysis of multiple pathogenic isolates of Streptococcus agalactiae: Implications for the microbial “pan-genome.” Proc. Natl. Acad. Sci. 102, 13950–13955 (2005).

19. H. Tettelin, D. Riley, C. Cattuto, D. Medini, Comparative genomics: the bacterial pan-genome. Curr. Opin. Microbiol. 11, 472–477 (2008).

20. L. V. Mallet, J. Becq, P. Deschavanne, Whole genome evaluation of horizontal transfers in the pathogenic fungus Aspergillus fumigatus. BMC Genomics 11, 171 (2010).

21. M. Marcet-Houben, T. Gabaldón, Acquisition of prokaryotic genes by fungal genomes. Trends Genet. TIG 26, 5–8 (2010).

22. M. Nguyen, A. Ekstrom, X. Li, Y. Yin, HGT-Finder: A New Tool for Horizontal Gene Transfer Finding and Application to Aspergillus genomes. Toxins 7, 4035–4053 (2015).

23. A. S. Urquhart, S. O’Donnell, E. Gluck-Thaler, A. A. Vogan, A natural mechanism of eukaryotic horizontal gene transfer. [Preprint] (2025). Available at: https://www.biorxiv.org/content/10.1101/2025.02.28.640899v1 [Accessed 14 April 2025].

24. A. Urquhart, A. A. Vogan, E. Gluck-Thaler, Starships: a new frontier for fungal biology. Trends Genet. TIG 40, 1060–1073 (2024).

25. E. Gluck-Thaler, et al., Giant transposons promote strain heterogeneity in a major fungal pathogen. mBio 16, e01092–25 (2025).

26. B. Auxier, et al., The human fungal pathogen Aspergillus fumigatus can produce the highest known number of meiotic crossovers. PLOS Biol. 21, e3002278 (2023).

27. G. Tonkin-Hill, et al., Robust analysis of prokaryotic pangenome gene gain and loss rates with Panstripe. Genome Res. 33, 129–140 (2023).

28. A. S. Urquhart, A. A. Vogan, D. M. Gardiner, A. Idnurm, Starships are active eukaryotic transposable elements mobilized by a new family of tyrosine recombinases. Proc. Natl. Acad. Sci. 120, e2214521120 (2023).

29. E. Gluck-Thaler, et al., Giant Starship Elements Mobilize Accessory Genes in Fungal Genomes. Mol. Biol. Evol. 39, msac109 (2022).

30. N. J. Croucher, et al., Diversification of bacterial genome content through distinct mechanisms over different timescales. Nat. Commun. 5, 5471 (2014).

31. S. A. Fritz, A. Purvis, Selectivity in Mammalian Extinction Risk and Threat Types: a New Measure of Phylogenetic Signal Strength in Binary Traits. Conserv. Biol. 24, 1042–1051 (2010).

32. S. Manzano-Morales, Y. Liu, S. González-Bodí, J. Huerta-Cepas, J. Iranzo, Comparison of gene clustering criteria reveals intrinsic uncertainty in pangenome analyses. Genome Biol. 24, 250 (2023).

33. G. Tonkin-Hill, J. Corander, J. Parkhill, Challenges in prokaryote pangenomics. Microb. Genomics 9, mgen001021 (2023).

34. W. Du, et al., The C2H2 Transcription Factor SltA Contributes to Azole Resistance by Coregulating the Expression of the Drug Target Erg11A and the Drug Efflux Pump Mdr1 in Aspergillus fumigatus. Antimicrob. Agents Chemother. 65, 10.1128/aac.01839-20 (2021).

35. J. W. Slaven, et al., Increased expression of a novel Aspergillus fumigatus ABC transporter gene, atrF, in the presence of itraconazole in an itraconazole resistant clinical isolate. Fungal Genet. Biol. FG B 36, 199–206 (2002).

36. D. Hagiwara, et al., A Novel Zn2-Cys6 Transcription Factor AtrR Plays a Key Role in an Azole Resistance Mechanism of Aspergillus fumigatus by Co-regulating cyp51A and cdr1B Expressions. PLOS Pathog. 13, e1006096 (2017).

37. M. G. Fraczek, et al., The cdr1B efflux transporter is associated with non-cyp51a-mediated itraconazole resistance in Aspergillus fumigatus. J. Antimicrob. Chemother. 68, 1486–1496 (2013).

38. S. Chen, et al., Variability in competitive fitness among environmental and clinical azole-resistant Aspergillus fumigatus isolates. mBio 15, e00263–24 (2024).

39. E. Mavridou, et al., Composite Survival Index to Compare Virulence Changes in Azole-Resistant Aspergillus fumigatus Clinical Isolates. PLoS ONE 8, e72280 (2013).

40. I. Valsecchi, E. Mellado, R. Beau, S. Raj, J.-P. Latgé, Fitness Studies of Azole-Resistant Strains of Aspergillus fumigatus. Antimicrob. Agents Chemother. 59, 7866–7869 (2015).

41. E. A. Hatmaker, et al., Population structure in a fungal human pathogen is potentially linked to pathogenicity. Nat. Commun. 16, 7594 (2025).

42. F. Baquero, T. M. Coque, J. C. Galán, J. L. Martinez, The Origin of Niches and Species in the Bacterial World. Front. Microbiol. 12, 657986 (2021).

43. M. A. Brockhurst, et al., The Ecology and Evolution of Pangenomes. Curr. Biol. CB 29, R1094–R1103 (2019).

44. M. Touchon, et al., Phylogenetic background and habitat drive the genetic diversification of Escherichia coli. PLoS Genet. 16, e1008866 (2020).

45. R. J. Hall, et al., Gene-gene relationships in an Escherichia coli accessory genome are linked to function and mobility. Microb. Genomics 7, 000650 (2021).

46. M. C. Fisher, et al., Tackling the emerging threat of antifungal resistance to human health. Nat. Rev. Microbiol. 20, 557–571 (2022).

47. A. Bankevich, et al., SPAdes: A New Genome Assembly Algorithm and Its Applications to Single-Cell Sequencing. J. Comput. Biol. 19, 455–477 (2012).

48. P. Bowyer, A. Currin, D. Delneri, M. G. Fraczek, Telomere-to-telomere genome sequence of the model mould pathogen Aspergillus fumigatus. Nat. Commun. 13, 5394 (2022).

49. W. C. Nierman, et al., Genomic sequence of the pathogenic and allergenic filamentous fungus Aspergillus fumigatus. Nature 438, 1151–1156 (2005).

50. D. E. Wood, J. Lu, B. Langmead, Improved metagenomic analysis with Kraken 2. Genome Biol. 20, 257 (2019).

51. F. A. Simão, R. M. Waterhouse, P. Ioannidis, E. V. Kriventseva, E. M. Zdobnov, BUSCO: assessing genome assembly and annotation completeness with single-copy orthologs. Bioinformatics 31, 3210–3212 (2015).

52. G. S. C. Slater, E. Birney, Automated generation of heuristics for biological sequence comparison. BMC Bioinformatics 6, 31 (2005).

53. L. Gabriel, et al., BRAKER3: Fully automated genome annotation using RNA-seq and protein evidence with GeneMark-ETP, AUGUSTUS and TSEBRA. BioRxiv Prepr. Serv. Biol. 2023.06.10.544449 (2024). 10.1101/2023.06.10.544449.

54. D. Kuznetsov, et al., OrthoDB v11: annotation of orthologs in the widest sampling of organismal diversity. Nucleic Acids Res. 51, D445–D451 (2022).

55. B. Buchfink, C. Xie, D. H. Huson, Fast and sensitive protein alignment using DIAMOND. Nat. Methods 12, 59–60 (2015).

56. R. J. Dutton, T. Reiter, PreHGT: A scalable workflow that screens for horizontal gene transfer within and between kingdoms. Arcadia Sci. (2023). 10.57844/arcadia-jfbp-7p11.

57. T. Reiter, Clustering the NCBI nr database to reduce database size and enable faster BLAST searches. Arcadia Sci. (2023).

58. D. M. Emms, S. Kelly, OrthoFinder: phylogenetic orthology inference for comparative genomics. Genome Biol. 20, 238 (2019).

59. R Core Team, R: A Language and Environment for Statistical Computing (R Foundation for Statistical Computing, 2021).

60. E. Paradis, K. Schliep, ape 5.0: an environment for modern phylogenetics and evolutionary analyses in R. Bioinforma. Oxf. Engl. 35, 526–528 (2019).

61. G. Yu, D. K. Smith, H. Zhu, Y. Guan, T. T.-Y. Lam, ggtree: an r package for visualization and annotation of phylogenetic trees with their covariates and other associated data. Methods Ecol. Evol. 8, 28–36 (2017).

62. G. Yu, T. T.-Y. Lam, H. Zhu, Y. Guan, Two Methods for Mapping and Visualizing Associated Data on Phylogeny Using Ggtree. Mol. Biol. Evol. 35, 3041–3043 (2018).

63. D. Hernangómez, giscoR: Download Map Data from GISCO API - Eurostat (2026).

64. E. Pebesma, Simple Features for R: Standardized Support for Spatial Vector Data. R J. 10, 439–446 (2018).

65. E. Pebesma, R. Bivand, Spatial Data Science: With applications in R (Chapman and Hall/CRC, 2023).

66. G. Yu, scatterpie: Scatter Pie Plot (2025).

67. H. Wickham, ggplot2: Elegant Graphics for Data Analysis (Springer-Verlag New York, 2016).

68. H. Chown, retroPal. (2026). https://doi.org/NA. Deposited February 2026.

69. H. Wickham, R. François, L. Henry, K. Müller, D. Vaughan, dplyr: A Grammar of Data Manipulation (2023).

70. H. Wickham, D. Vaughan, M. Girlich, tidyr: Tidy Messy Data (2025).

71. C. Camacho, et al., BLAST+: architecture and applications. BMC Bioinformatics 10, 421 (2009).

72. G. Van Rossum, F. L. Drake, Python 3 Reference Manual (CreateSpace, 2009).

73. A. Hagberg, P. J. Swart, D. A. Schult, “Exploring network structure, dynamics, and function using NetworkX” (Los Alamos National Laboratory (LANL), Los Alamos, NM (United States), 2008).

74. P. Jones, et al., InterProScan 5: genome-scale protein function classification. Bioinformatics 30, 1236–1240 (2014).

75. O. Brynildsrud, J. Bohlin, L. Scheffer, V. Eldholm, Rapid scoring of genes in microbial pan-genome-wide association studies with Scoary. Genome Biol. 17, 238 (2016).

76. L. Snipen, K. H. Liland, micropan: an R-package for microbial pan-genomics. BMC Bioinformatics 16, 79 (2015).

77. R. Bouckaert, et al., BEAST 2.5: An advanced software platform for Bayesian evolutionary analysis. PLOS Comput. Biol. 15, e1006650 (2019).

78. D. Orme, et al., caper: Comparative Analyses of Phylogenetics and Evolution in R (2025).

79. S. N. Wood, Generalized Additive Models: An Introduction with R, Second Edition, 2nd Ed. (Chapman and Hall/CRC, 2017).

80. S. N. Wood, Stable and Efficient Multiple Smoothing Parameter Estimation for Generalized Additive Models. J. Am. Stat. Assoc. 99, 673–686 (2004).

81. S. Coretta, tidymv: Tidy model visualisation for generalised additive models. R Package Version 3 (2022).

82. P. O. Lewis, A Likelihood Approach to Estimating Phylogeny from Discrete Morphological Character Data. Syst. Biol. 50, 913–925 (2001).

83. L. J. Revell, phytools 2.0: an updated R ecosystem for phylogenetic comparative methods (and other things). PeerJ 12, e16505 (2024).

84. F. J. Whelan, M. Rusilowicz, J. O. McInerney, Coinfinder: detecting significant associations and dissociations in pangenomes. Microb. Genomics 6, e000338 (2020).

85. G. Csárdi, et al., igraph: Network Analysis and Visualization in R (2025).

86. M. Bastian, S. Heymann, M. Jacomy, Gephi: An Open Source Software for Exploring and Manipulating Networks. Proc. Int. AAAI Conf. Web Soc. Media 3, 361–362 (2009).

87. E. Gluck-Thaler, A. A. Vogan, Systematic identification of cargo-mobilizing genetic elements reveals new dimensions of eukaryotic diversity. Nucleic Acids Res. 52, 5496–5513 (2024).

88. S. R. Eddy, Accelerated Profile HMM Searches. PLOS Comput. Biol. 7, 1–16 (2011).

