## Supplemental Figures and Info for "The pangenome of *Aspergillus fumigatus* highlights the dynamics of gene gain-loss over evolutionary timescales in a human fungal pathogen"

**This PDF file includes:**

Figures S1 to S3
Legends for Datasets S1 to S4
SI References

**Other supporting materials for this manuscript include the following:**

Datasets S1 to S4

### Figures

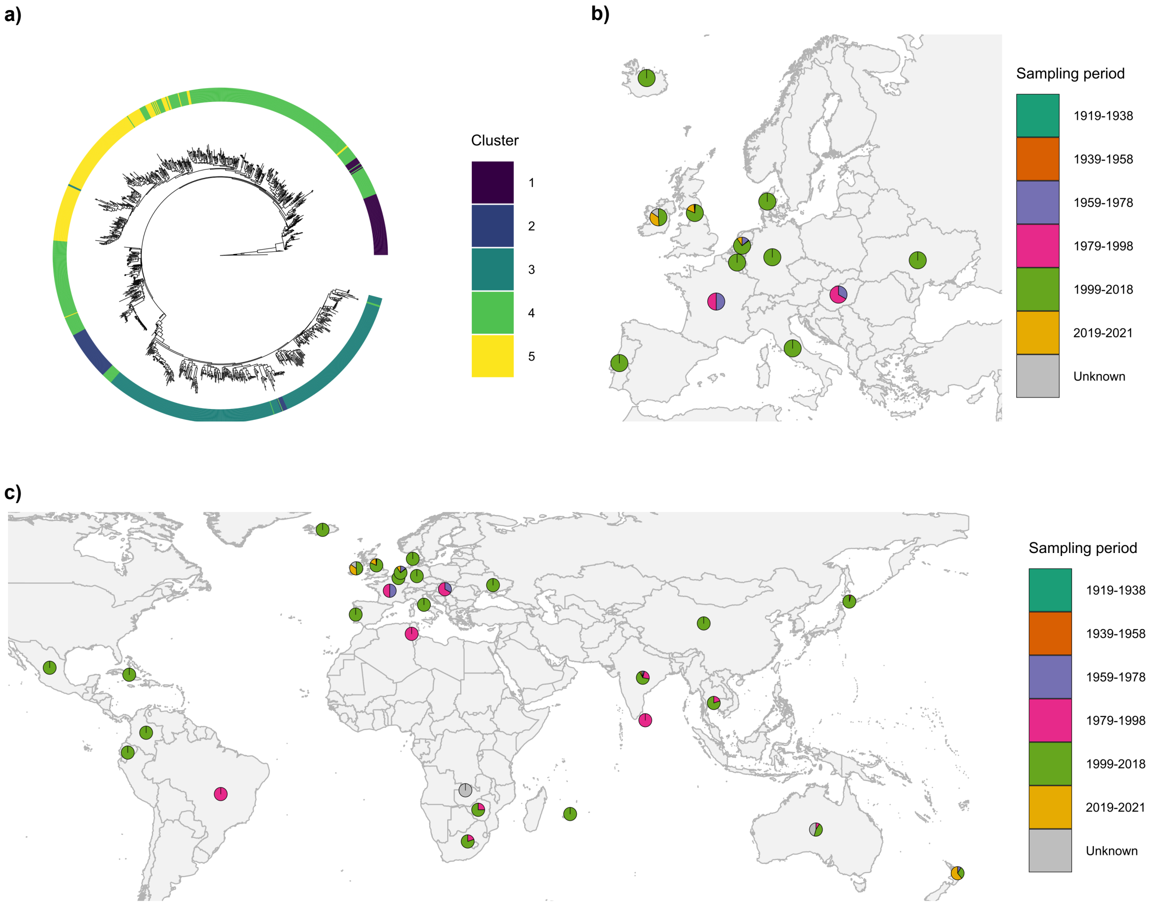


**Fig. S1. Phylogenetic and geographic distribution of underlying data. a)** Reconstruction of phylogenetic tree from M. Fisher et al., 2025 (1) depicting 1,098 isolates used in the pangenome of this study. Colours are based on assigned lineages from M. Fisher et al., 2025. **b-c)** European **(b)** and worldwide (**c**) distribution of isolates. Pie-charts depict the proportion of isolates from a range of sampling periods.

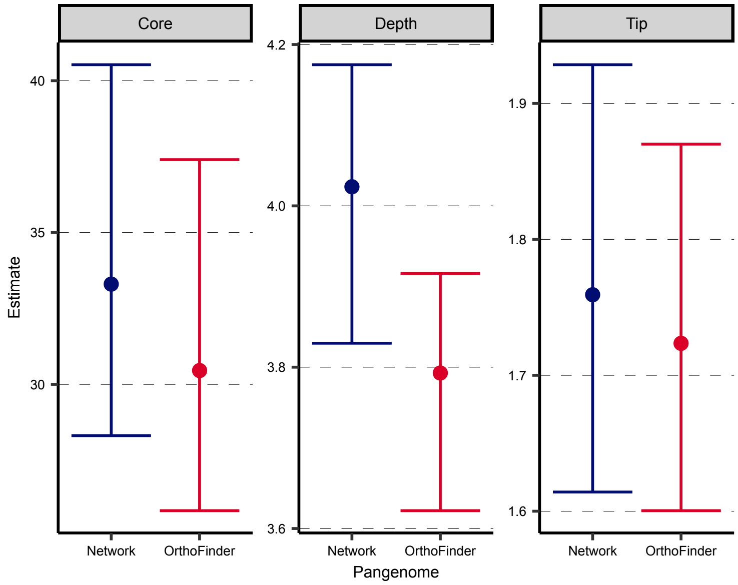


**Fig. S2. Pangenome statistics across orthogroup models**. Estimates of inferred coefficients from Panstripe fitted generalised linear models, using a SNP-based phylogeny. Core refers to whether branch lengths in the phylogeny are associated with orthogroup gain and loss. Depth indicates whether the rate of gene gain and loss changes significantly with the depth of the branch. Tip infers that associations with genes are observed to occur on the tips of the phylogeny. All estimates were significant ($p < 0.05$). Error bars represent the 25% and 75% confidence interval.

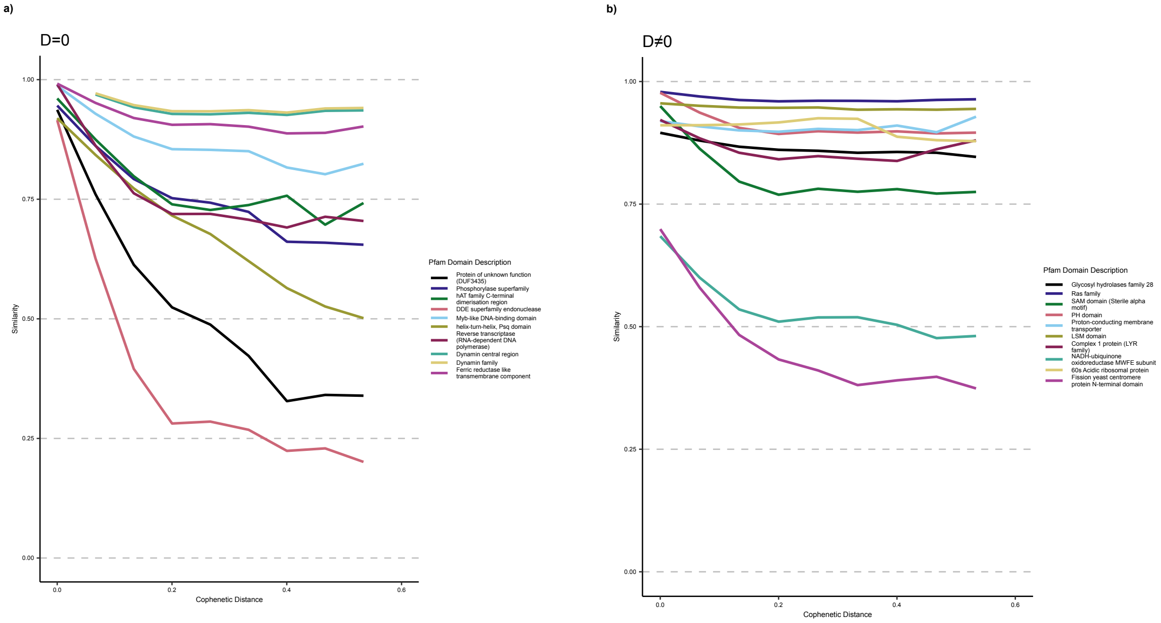


**Fig. S3. Gain and loss dynamics across lineage-dependent and lineage-independent orthogroups.** Generalised additive model (GAM) fits describing the relationship between cophenetic (pairwise phylogenetic) distance and Jaccard similarity of presence-absence profiles for selected orthogroups with known Pfam domains. **a)** Subset of lineage-dependent orthogroups containing Pfam domains **b)** Subset of lineage-independent orthogroups containing Pfam domains

**Dataset S1 (separate file).** Pangenome matrix from network clustering method.

**Dataset S2 (separate file).** InterProScan protein domains of representative pangenome using orthogroup hubs; panGWAS results from Scoary combined with InterProScan annotations.

**Dataset S3 (separate file).** *Starship* captain family assignment from hmmsearch.

**Dataset S4 (separate file).** D-statistic for each accessory orthogroup; edge-list of lineage-independent accessory orthogroups, from Coinfinder.

### SI References

1. M. Fisher, *et al.*, Recent European origin of azole resistance in the critical priority fungal pathogen Aspergillus fumigatus. [Preprint] (2025). Available at: https://www.researchsquare.com/article/rs-7905776/v1 [Accessed 9 March 2026].
